# EdU incorporation to assess cell proliferation and drug susceptibility for the brain-eating amoeba, *Naegleria fowleri*

**DOI:** 10.1101/2021.01.07.425827

**Authors:** Emma V. Troth, Dennis E. Kyle

**Affiliations:** Department of Infectious Diseases, University of Georgia, Athens, Georgia, USA; Center for Tropical and Emerging Global Diseases, Athens, Georgia, USA; Department of Cellular Biology, University of Georgia, Athens, Georgia, USA

## Abstract

*Naegleria fowleri* is a pathogenic free-living amoeba that is commonly found in warm, freshwater and can cause a rapidly fulminant disease known as primary amoebic meningoencephalitis (PAM). New drugs are urgently needed to treat PAM, as the fatality rate is >97%. Until recently, few advances have been made in the discovery of new drugs for *N. fowleri* and one drawback is the lack of validated tools and methods to enhance drug discovery and diagnostics research. In this study we aimed to validate alternative methods to assess cell proliferation that are commonly used for other cell types and develop a novel drug screening assay to evaluate drug efficacy on *N. fowleri* replication. EdU (5-ethynyl-2′-deoxyuridine) is a pyrimidine analog of thymidine that can be used as a quantitative endpoint for cell proliferation. EdU incorporation is detected via a copper catalyzed click reaction with an Alexa Fluor linked azide. EdU incorporation in replicating *N. fowleri* was validated using fluorescence microscopy and quantitative methods for assessing EdU incorporation were developed by using an imaging flow cytometer. Currently used PAM therapeutics inhibited *N. fowleri* replication and EdU incorporation *in vitro*. EdA (5′ethynyl-2′-deoxyadenosine), an adenine analog, also was incorporated by N. fowleri, but was more cytotoxic than EdU. In summary, EdU incorporation could be used as a complimentary method for drug discovery for these neglected pathogens.

## Introduction

*Naegleria fowleri* is a free-living pathogenic amoeba that is ubiquitous in freshwater and soil and poses a significant public health risk as the causative agent of Primary Amebic Meningoenchephalitis (PAM), a disease with a ≥ 97% fatality rate [1]. Individuals can be infected by intranasal instillation of amoebae in untreated or poorly treated water from natural bodies of water (lakes, ponds, rivers) or tap water and swimming pools with insufficient chlorination. Infection via tap water has historically been the result of Netipot use and religious practices, such as ritual ablution [2]. Once inside the nasal cavity, *N. fowleri* traverses the nasal epithelium and travels the length of the olfactory nerve through the cribriform plate, and enters the frontal lobe of the brain where it begins to degrade brain tissue. Death typically occurs within 5-12 days after infection. PAM is traditionally treated with a cocktail of antifungal and antibiotic drugs that usually includes amphotericin B, azithromycin, an azole (fluconazole or ketoconazole), and miltefosine [2]. The drug regimen was derived empirically and has changed very little since the early 1970s, underscoring how neglected drug discovery research is for this pathogen.

*Naegleria* is a genus of over 40 species, but only one species, *Naegleria fowleri*, has been confirmed to be pathogenic in humans [3]. *N. fowleri* has three life stages: trophozoite, cyst, and flagellate. The trophozoite form is the active, feeding stage of the amoeba; in this stage, the amoeba replicates via promitosis every 10-12 hours [4]. *N. fowleri* may transform into a dormant cyst stage in unfavorable environmental conditions. Of note, *N. fowleri* cysts have never been found in the brain of PAM patients[1]. The flagellated form enhances motility of *N. fowleri* in water or media, but this life stage of *N. fowleri* is the least well studied.

In the years since its discovery to be the causative agent of PAM in 1965, *N. fowleri* has received sparse research attention despite its significant mortality rate [5]. Until recently, little effort has been applied to the field of drug discovery for these amoebae. High-throughput screening assays have now been developed to identify novel compounds active against *N. fowleri* [6, 7] and screening of multiple compound repurposing libraries have yielded a number of promising compounds [8–11]. Though the development of a high-throughput screen has had a significant impact on the field of drug discovery for this pathogen, there is still a need for secondary assays to confirm these drug hits and to reveal information regarding the mechanism of action of these compounds.

To further investigate the biology of *N. fowleri* and to develop a secondary assay to evaluate the effect of compounds of *N. fowleri* replication, we investigated methods used to assess cell proliferation for other cells. 5-ethynyl-2′-deoxyuridine (EdU) is a nucleoside analog of thymidine that is incorporated into DNA during active cellular replication and detection utilizes click-chemistry. EdU contains an alkyne group that can undergo a copper catalyzed click reaction with the azide group on an AlexaFluor probe. EdU has been useful to assess cell proliferation in a wide variety of parasites, including *Cryptosporidium parvum, Schistosoma mansoni, Trypanosoma brucei, Trypanosoma cruzi*, and *Acanthamoeba castellanii* [12–15]. In parallel, the purine analog 7-Deaza-2′-deoxy-7-ethynyladenosine (EdA) has been used for similar purposes in mammalian cells [16].

In this study, EdU and EdA were assessed for their ability to incorporate into DNA of logarithmically growing trophozoites of *N. fowleri*. A standard protocol was developed and detection of EdU and EdA incorporation was assessed by imaging flow cytometry. The EdU assay was then used to assess potency of the current treatment PAM drugs azithromycin, amphotericin B, and miltefosine, along with a promising repurposed drug lead, posaconazole [9]. Fifty percent inhibitory (IC_50_) concentrations generated with the EdU assay were comparable to the standard CellTiter-Glo high-throughput screening assay [6]. In summary, we report the utility of EdU and EdA incorporation for assessing cell proliferation and drug screening for *N. fowleri*.

## Results

### Validation of EdU Incorporation Assay

The thymidine analog, EdU, was assessed for its ability to incorporate into *N. fowleri* trophozoite DNA during logarithmic growth *in vitro*. EdU is structurally identical to thymidine aside from an alkyne handle modification (Figure S1). EdU incorporation was detected via a copper-catalyzed alkyne-azide click reaction with an azide containing AlexaFluor probe. Once EdU incorporates into DNA, the AlexaFluor azide will click to it in the presence of copper and result in a fluorescent probe. For these studies we used the manufacturer recommended EdU concentration of 10 μM during labelling studies. The EdU detection protocol was slightly modified from the manufacturer’s recommendation (Figure S2) by using 1% saponin instead 0.5% Triton-X for cell permeabilization; this process resulted in better preservation of the amoeboid shape of *N. fowleri* trophozoites (data not shown).

We assessed EdU as a measure of cell replication by incubating *N. fowleri* with 10 μM EdU for 72 hours. The EdU detection protocol was performed and labelled *N. fowleri* were mounted on glass microscope slides. Upon visualization with high-resolution fluorescence microscopy, *N. fowleri* efficiently incorporated EdU with no observed cytotoxicity (Figure 1). EdU staining colocalized with Hoechst nuclear stain confirming EdU localization in the nucleus.

**Figure 1.**
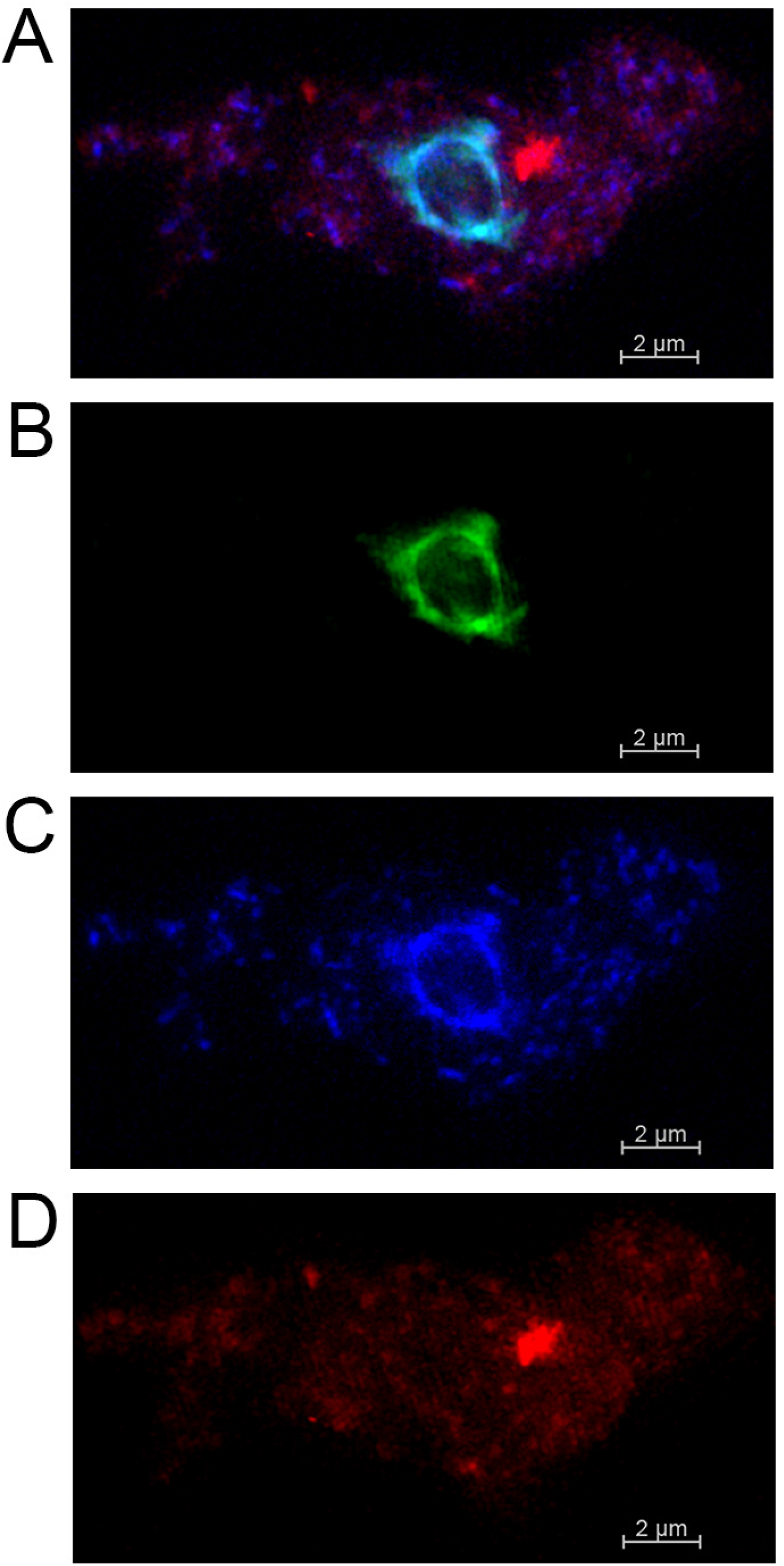
EdU incorporation in replicating *N. fowleri*. Detection of EdU in *N. fowleri* trophozoites was assessed after 72-hour incubation with 10 μM EdU. Images were obtained on Zeiss ELYRA S1 microscope at 100x. **A.** Merged channels image of EdU (green), Hoescht (blue), and Cytopainter Red (red). B. EdU staining. **C.** 1 μg/mL Hoescht staining. **D.** 20x Cytopainter Red staining.

### Detection of EdU Incorporation by Imaging Flow Cytometry

Most cell proliferation methods currently used for *N. fowleri* are unable to quantify amoeba number and simultaneously collect morphological data in a high-throughput manner. Therefore, a potential advantage of EdU is the ability to assess incorporation by imaging flow cytometry. We used an ImageStream (Millipore) flow cytometer to develop the EdU analysis protocol because it allows quantification of cell numbers as well as capture of an image of every cell that passes through the cytometer.

Imaging flow cytometry potentially gives us the ability to evaluate the morphological phenotype of *N. fowleri* trophozoites as well as quantifying EdU incorporation.

For the quantitative imaging assay, we first determined the gating parameters. The average size of a *N. fowleri* trophozoite is 15 – 25 μm [4]. After exposure of *N. fowleri* trophozoites to amphotericin B, azithromycin, miltefosine, or posaconazole for 72 hours, we found a minimum diameter gate of 7.5 μM captured intact *N. fowleri* trophozoites after drug treatment (data not shown). This 7.5 μM diameter was the smallest diameter that captured intact and potentially viable *N. fowleri* trophozoites and excluded cell debris.

We next assessed the optimal timing for addition of EdU that would support development of a drug susceptibility assay. The current standard drug assay is 72 hours and is designed to identify fast acting compounds required to advance therapy of PAM [6, 8–11]. Initially we added EdU at 48 hours and then incubated the cells for another 24 hours and found 50-60% of *N. fowleri* trophozoites were EdU positive (Figure 2). When EdU was added at time 0 and incubated for 72 hours with amoebae, ≥90% of *N. fowleri* trophozoites were EdU positive.

**Figure 2.**
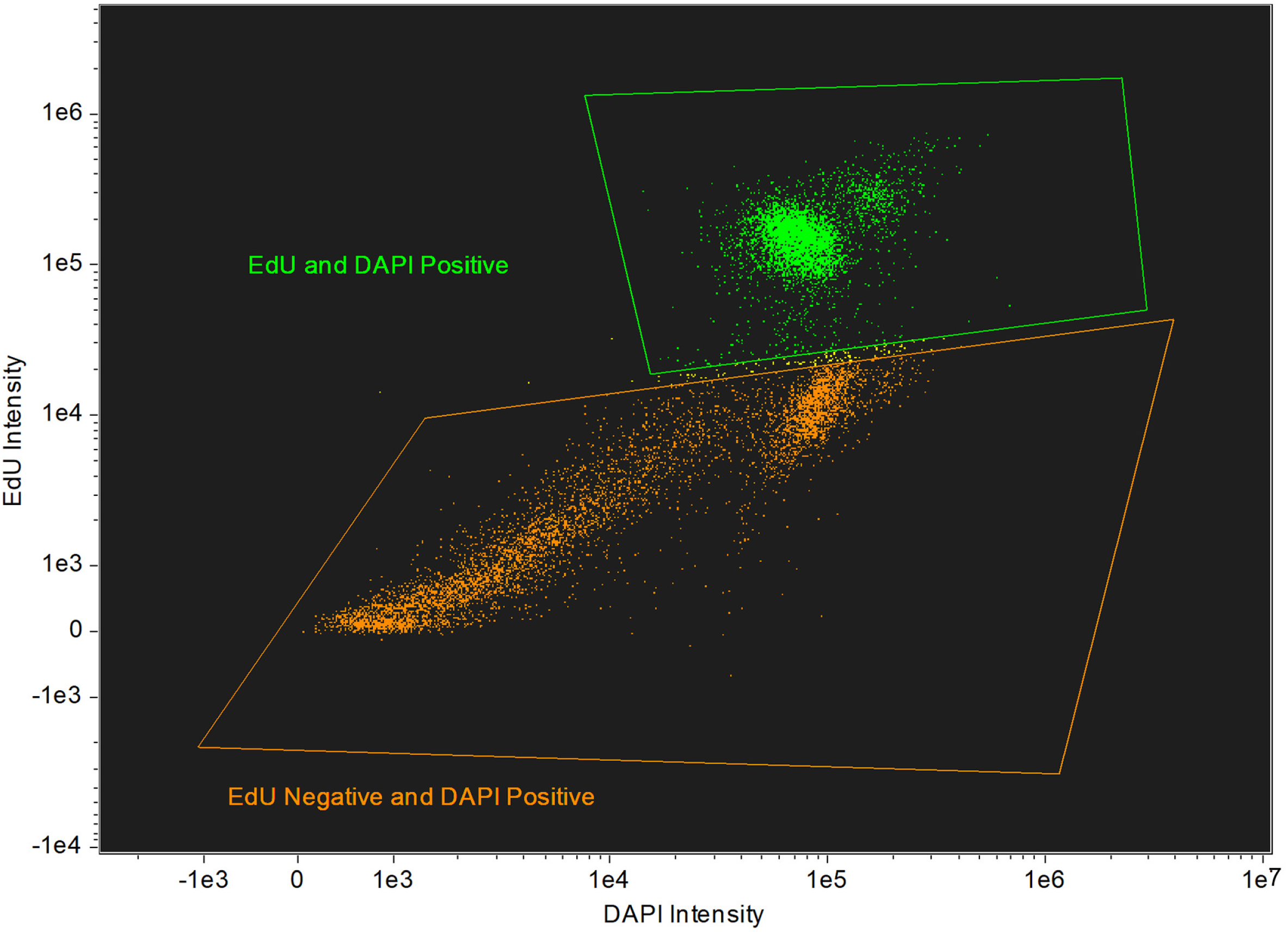
Populations of EdU positive and EdU negative *N. fowleri* after EdU addition at 48 hours. 20,000 *N. fowleri* trophozoites were incubated in 500 μL NCM for 72 hours. 10 μM EdU was added at 48 hours. Cells were fixed, permeabilized, stained, and analyzed on the Millipore ImageStream. Approximately 55% of the 10,000 captured cells were EdU positive.

### EdU Drug Susceptibility Assay

Amphotericin B, azithromycin, miltefosine, and posaconazole are standard drugs for treating PAM and they were used for evaluation of the EdU drug susceptibility assay. *N. fowleri* trophozoites were exposed to serially diluted concentrations of each drug for 72 hours. EdU was added at 48 hours and the percent of EdU positive cells in untreated controls was used to normalize the data. The imaging flow cytometry data acquisition template was set to capture 10,000 events per sample. Flow cytometry data was compensated and analyzed using the IDEAS software (Luminex). Single color, as well as unstained controls were included with each replicate; percent EdU positive cells were normalized to the untreated controls. The imaging cytometry data was used to generate concentration response data and to calculate IC_50s_ for the four compounds tested (Figure 3). The IC_50_ values determined for amphotericin B, azithromycin, miltefosine, posaconazole were 0.027 μM, 0.112 μM, 47.77 μM, 0.002 μM and respectively.

**Figure 3.**
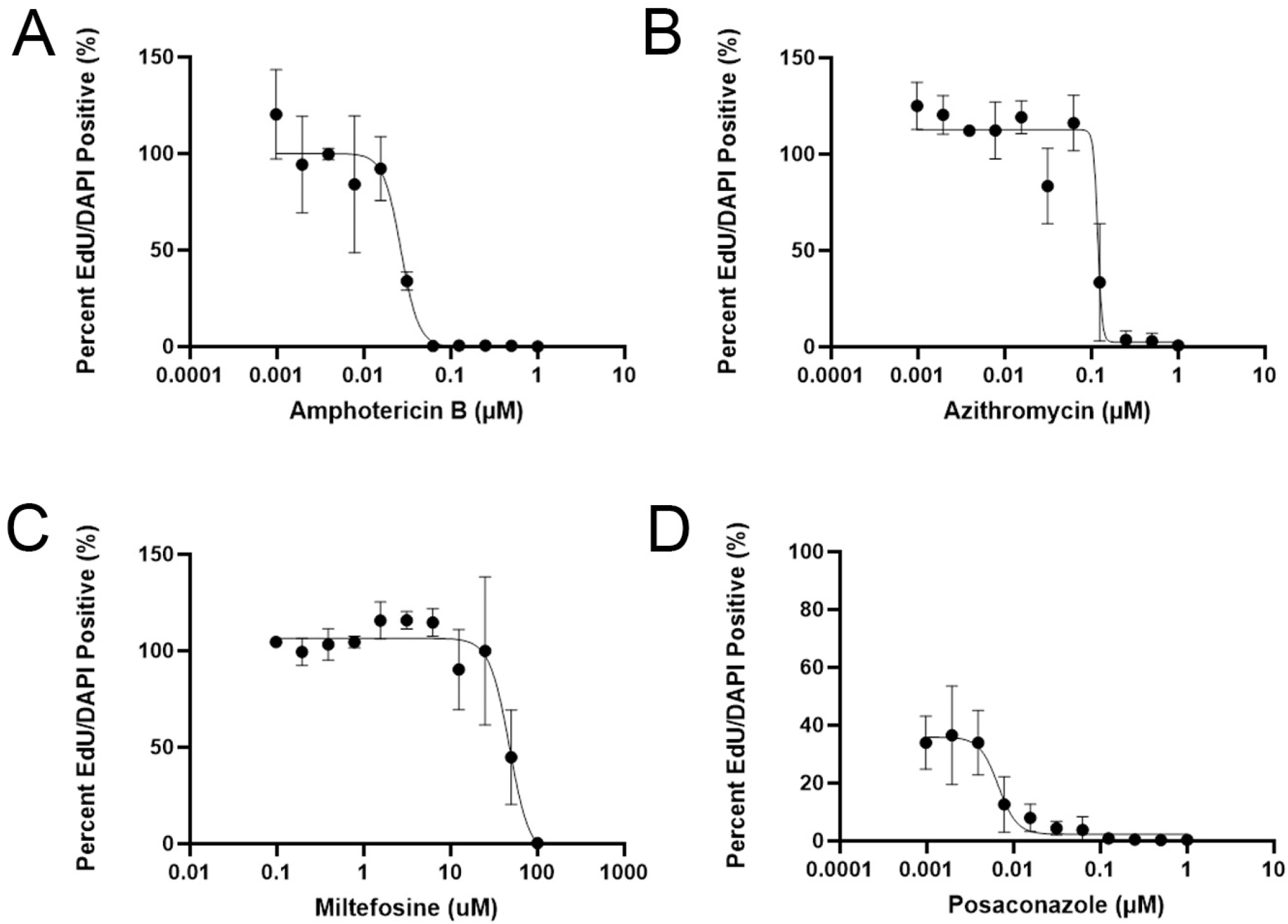
IC_50_ Curves generated by the EdU assay. *N. fowleri* trophozoites were incubated with serial dilutions of amphotericin B (**A**), azithromycin (**B**), miltefosine (**C**), and posaconazole (**D**) for 72 hours. 10 μM EdU was added at 48 hours. Cells were fixed, permeabilized, and stained at 72 hours. Cells were analyzed on the Millipore ImageStream flow cytometer. Unstained and single-color controls were included for each replicate. IC_50_ values were determined to be: 0.027 μM (amphotericin B), 0.112 μM (azithromycin), 0.002 μM (posaconazole), 47.77 μM (miltefosine) (*n = 2*).

Once we established the EdU assay could be used to generate drug concentration responses, we compared the IC_50_ values obtained from the EdU assay, which measures cell replication, to the standard CellTiter-Glo assay used for high throughput screening. CellTiter-Glo assesses metabolic activity (ATP content). Amphotericin B, azithromycin, miltefosine, and posaconazole were serially diluted and screened in the 72-hour CellTiter-Glo assay with *N. fowleri* trophozoites. The IC_50s_ generated in the two assays for the four drugs tested fall within a 10-fold range (Table 1).

**Table 1.** Comparison of IC_50_ values generated by the EdU assay and the CellTiter-Glo assay. IC_50_ values were generated by either the EdU or CellTiter-Glo assays. For both assays, *N. fowleri* trophozoites were exposed to drug for a total of 72 hours.

|  | EdU IC<sub>50</sub> (μM) | CellTiter-Glo IC<sub>50</sub> (μM) |
| --- | --- | --- |
| Amphotericin B | 0.027 ± 0.01 | 0.04 |
| Azithromycin | 0.112 ± 0.05 | 0.01 |
| Miltefosine | 47.77 ± 11.6 | 60.5 ± 5.5 |
| Posaconazole | 0.002 ± 0.004 | 0.02 ± 0.01 |

As previously mentioned, imaging cytometry gives us the ability to evaluate the morphological phenotype of individual *N. fowleri* trophozoites as they pass through the cytometer. Drug-treated *N. fowleri* trophozoites were observed to be smaller and deformed compared to untreated *N. fowleri* (Figure 4). At the highest concentration of drugs tested, the vast majority of cells were EdU negative but remained Hoechst positive, suggesting that DNA replication was inhibited by drug.

**Figure 4.**
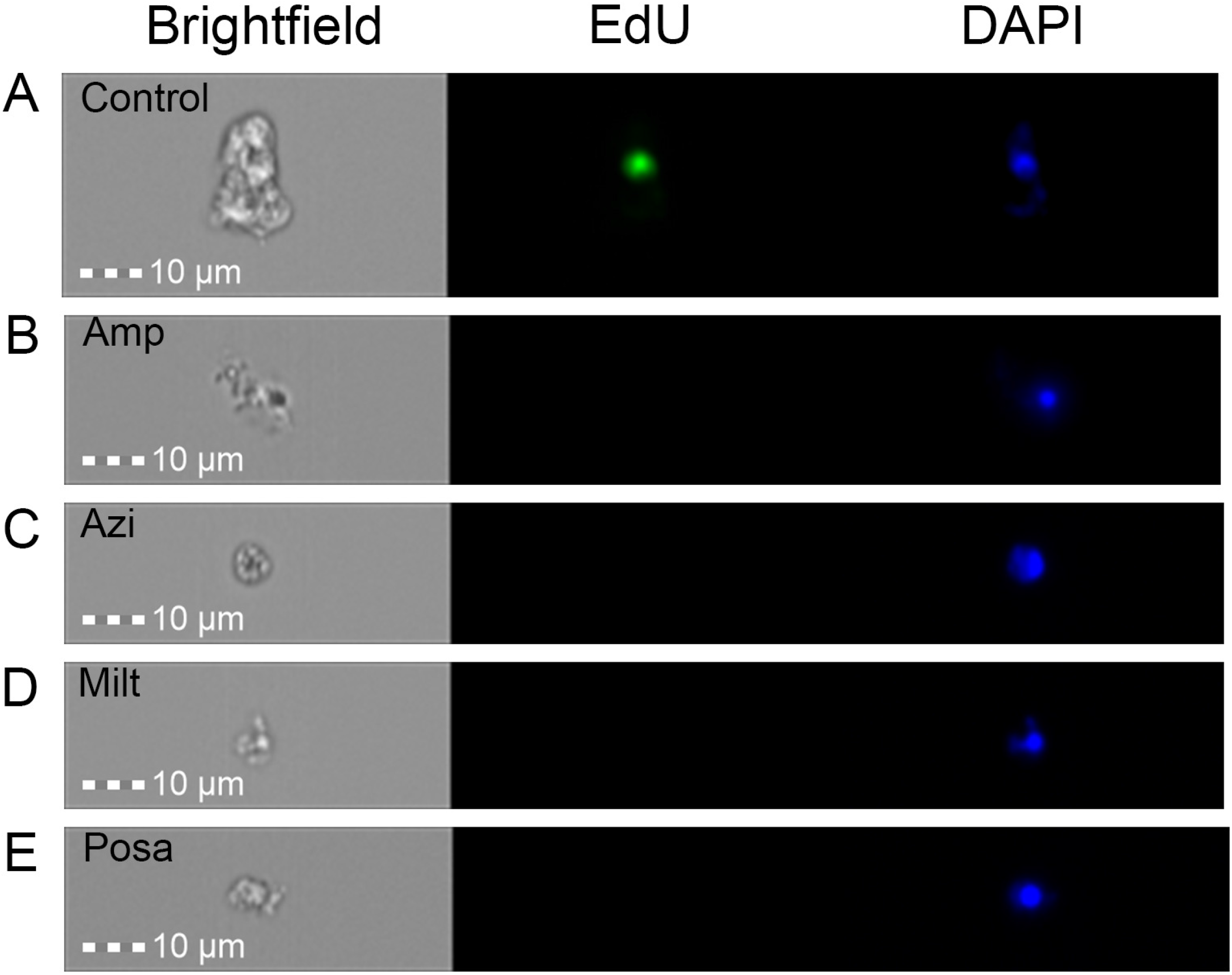
Phenotypic analysis of drug-treated *N. fowleri* trophozoites. Images of drug-treated *N. fowleri* trophozoites after incubation for 72 hr with **A)** 1 μM amphotericin B, **B)** 1 μM azithromycin, **C)** 100 μM miltefosine, or **D)** 1 μM Posaconazole. EdU (10 μM) added at 48 hours. Cells were fixed, permeabilized, and stained with EdU reaction mixture and 1 μg/mL Hoechst after 72 hours. Images were obtained from the Millipore ImageStream (40x).

### EdA incorporation in *N. fowleri* trophozoites

Although thymidine analogs are most commonly used for cell proliferation studies in most cell types, modified purines also can be used [16]. The adenosine analog EdA also contains an alkyne that is click-able with the AlexaFluor azide (Figure S1) and was evaluated for its ability to incorporate into *N. fowleri* trophozoite DNA during cell replication. EdA was observed to incorporate into *N. fowleri* DNA and colocalize with the nucleus based on Hoechst staining (Figure 5). Localization did not appear to be limited to the nucleus, though the area where Hoechst staining is present is the area of most intense EdA staining. It is possible the EdA is being incorporated into RNA as well as DNA.

**Figure 5.**
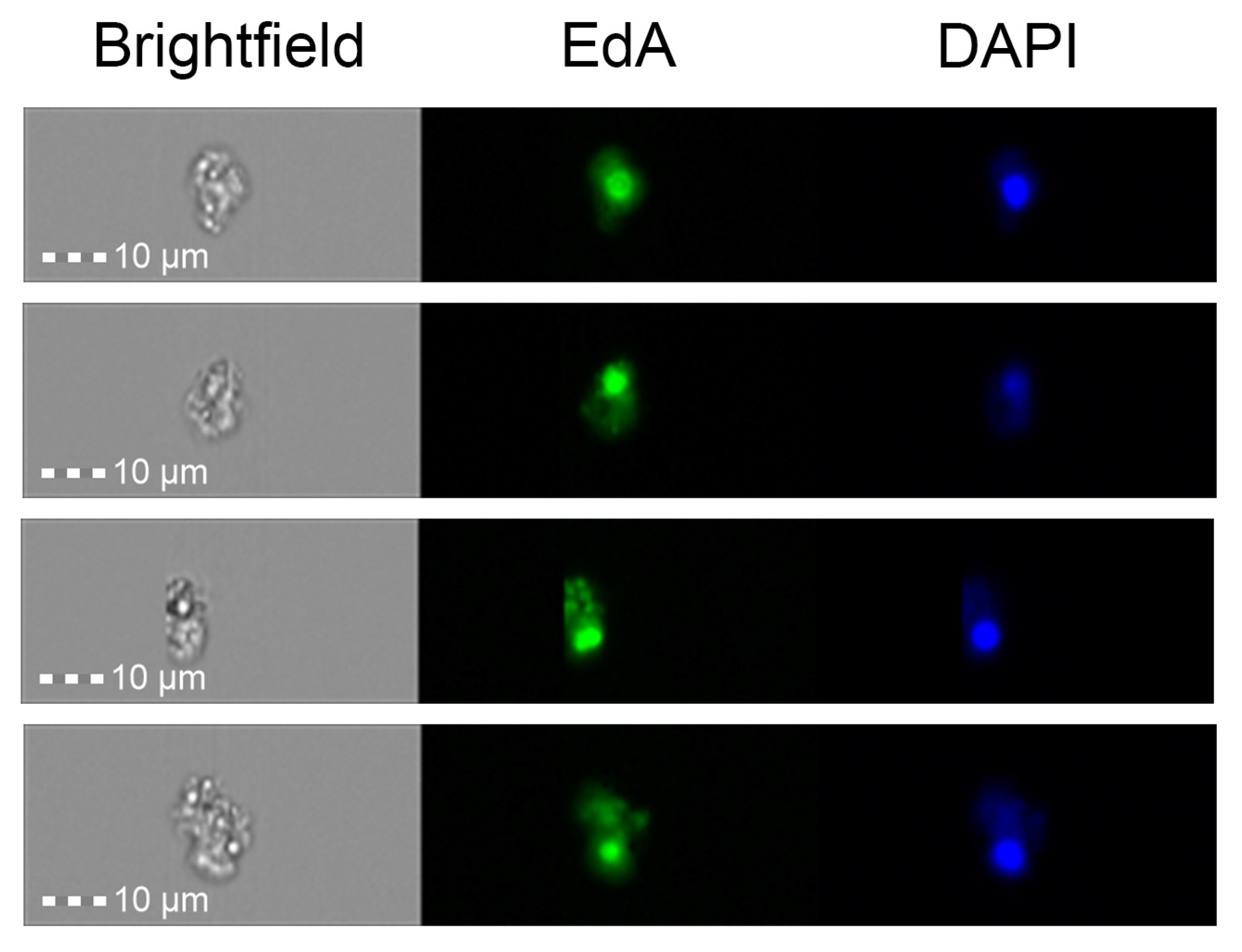
EdA incorporation in replicating *N. fowleri* trophozoites. Images of *N. fowleri* trophozoites after incubation with 10 μM EdA for 72 hours. Cells were fixed, permeabilized, and stained with EdU reaction mixture and 1 μg/mL Hoechst after 72 hours. Images were obtained from the Millipore ImageStream (40x).

Interestingly, EdA did not incorporate as efficiently as EdU in the 72 hr assay period. While ≥ 90% of cells were EdU positive after 72 hr incubation with 10 μM EdU, only 8.4 ± 3.8 % of cells were EdA positive after 72 hr incubation with 10 μM EdA (Figure S3). For these studies we used the same protocol of counting 10,000 cells within 5 minutes. These conditions were sufficient to routinely collect data for EdU positive cells in less than 5 minutes, but in the EdA experiments we only captured 1,968 ± 853 cells stained with EdA. These data suggest that 10 μM EdA is more cytotoxic to amoebae than equivalent concentrations of EdU.

## Discussion

This body of work documents the first report of the use of EdU to assess replication in *N. fowleri*. EdU efficiently incorporated in *N. fowleri* with no observed cytotoxicity. EdU staining colocalized exclusively with Hoechst nuclear staining, indicating EdU incorporation into DNA. Once EdU was incorporated into DNA, apparently it was detected even in dying cells (unless the integrity of the DNA is compromised during cell death). The EdU assay represents a new method for quantifying *N. fowleri* replication.

We developed an assay using EdU to assess the effects of drug exposure on *N. fowleri* replication. The current standard for drug discovery for *N. fowleri* is the CellTiter-Glo assay. CellTiter-Glo measures ATP content of the cell and can be categorized as a metabolic assay. Herein we have shown the EdU assay also can be used for drug discovery for *N. fowleri*. The current PAM therapeutics, amphotericin B, azithromycin, miltefosine, and posaconazole were evaluated in the EdU assay and were found to inhibit *N. fowleri* trophozoite replication at similar concentrations as the CellTiter-Glo assay. The EdU assay provides a novel way to assess drug efficacy by measuring cell replication. In addition to quantifiable drug concentration response data, the phenotypic effect of compounds on *N. fowleri* also were able to be assessed with the EdU assay. This assay was developed using the Millipore ImageStream flow cytometer but is adaptable to flow cytometers that do not have the ability to capture images of cell morphologies.

It is important to consider a few key similarities and differences between the EdU and the CellTiter-Glo assays for drug discovery. The CellTiter-Glo assay is compatible in 96- and 384-well plates. The EdU assay is most accurate and best suited to be performed in microcentrifuge tubes, thus resulting in lower throughput. The CellTiter-Glo assay requires 4,000 *N. fowleri* per well while the EdU assay requires 20,000 *N. fowleri* per tube. The volume of media and drug that *N. fowleri* grow in over the course of 72 hours is 100 μL in the CellTiter-Glo assay and 500 μL in the EdU assay. Some degree of variation is expected when comparing data from these assays; these include a morphological versus metabolic endpoint as well as different concentrations of amoebae used in the assay. Despite these differences between these assays, the IC_50_ values generated from the assays were comparable and fell within a 10-fold range. Although we assessed drugs with differing mechanisms of action, the similarities in assay results may differ for other mechanisms of action.

The major focus of this report has been on the use of EdU for *N. fowleri* drug discovery, but there are many broader implications as well. Currently, a lack of genetic tools available for use in *N. fowleri* limits almost all studies in this field. The use of EdU in *N. fowleri* can serve as another tool to provide insight in to the biology of these amoebae. For example, EdU incorporation can serve as a tool to examine DNA replication in response to a variety of environmental factors. Upon development of genetic tools such as gene knockout, molecular knockdown, and molecular tagging, EdU incorporation can be used to evaluate how various molecules within *N. fowleri* interact and effect cellular replication. EdU could also be used as a tool to examine the transition of *N. fowleri* between different life phases; trophozoite to cyst or vice versa. A similar study has been conducted in *Entamoeba invadens* [17]. EdU also has the advantage of not being radioactive. In *Acanthamoeba castellani*, EdU has been utilized to investigate the effect of one of its endosymbiotic bacteria, *Legionella pneumophilia*, on the amoeba’s ability to replicate [15]. EdU has been used extensively in other parasites for both drug discovery investigation as well as cellular biology studies. A stage-specific drug activity assay has been developed for *Cryptosporidium parvum* using EdU incorporation [12].

EdA also was evaluated for its ability to incorporate in to *N. fowleri* DNA. Though it was found to incorporate, the efficiency of incorporation was dramatically less than EdU and EdA was more cytotoxic to the amoebae. EdA is not recommended for drug susceptibility assays, yet it can serve other purposes. It is possible that EdA incorporates in to RNA as well as DNA, as it is a purine and images captured in this study showed EdA staining not being limited to the nucleus. Further investigation is warranted to determine if EdA incorporates into RNA.

The goal of this study was to identify whether EdU can incorporate in to replicating *N. fowleri* and if the EdU assay could be developed as a tool for drug discovery in these amoebae. Our results demonstrate that EdU incorporates into *N. fowleri* DNA and can be robustly detected. Furthermore, an EdU assay was developed to assess the effect of current treatment PAM therapeutics on *N. fowleri* replication. The EdU assay can provide novel insight into the mechanism of action of both current treatment therapeutics for PAM as well as lead compounds discovered from high-throughput screening campaigns. Performing this assay on the Millipore ImageStream flow cytometer also allows phenotypic assessment of drug compounds of *N. fowleri*. The EdU assay provides a secondary drug screening option for compounds identified in high-throughput screens. Screening molecules on this assay can provide insight in to their ability in inhibit cell replication and to rule out any false-positives that could have been generated in the CellTiter-Glo assay. The EdU assay offers a robust complementary drug discovery assay to aid in the identification of potent molecules for the treatment of PAM.

## Materials and Methods

### *Naegleria fowleri* culture

The *N. fowleri* strain (NF69) used in these studies was isolated from a fatal PAM case in 1969, a 9-year-old boy in Adelaide, Australia (ATCC 30215). Trophozoites were grown axenically at 34°C in Nelson’s Complete Media (NCM) supplemented with 10% fetal bovine serum (FBS) and 125 μg penicillin-streptomycin. All experiments were performed using logarithmic-phase trophozoites. All reagents were obtained from Sigma-Aldrich (St. Louis, MO).

### Evaluation of EdU incorporation

Presently, *N. fowleri* drug susceptibility assays reported in the literature evaluate drug efficacy based on metabolic properties, such as ATP content. Therefore, we assessed the potential to evaluate drug efficacy based on a novel parameter to provide further insight in to the ability of compounds to inhibit amoebic replication. The Click-iT Plus EdU Cell Proliferation Kit (Thermo Fisher Scientific, Waltham, MA) has not been used previously for *N. fowleri*. EdU (5′-ethynyl-2′-deoxyuridine) is a thymidine analog that can be incorporated in to DNA during active DNA replication.

To assess the ability of logarithmic-phase *N. fowleri* trophozoites to incorporate the EdU molecule, 20,000 trophozoites were seeded with 500 μL NCM and 10 μM EdU in Eppendorf 1.5 mL centrifuge tubes for 72 hours and incubated at 34°C. At 72 hours, the manufacturer’s protocol for EdU detection was followed with minor modifications. The Zeiss ELYRA S1 microscope does not have a bright-field lens; thus, a cytoplasmic stain was chosen to define the area of trophozoites and provide a reference for interpretation of EdU staining. Prior to fixation, *N. fowleri* were incubated with 10x AbCam Cytopainter Red for 1 hour at 34°C. Trophozoites were fixed with 2% paraformaldehyde in PBS for 20 minutes at room temperature. To remove liquid after each step of the protocol, Eppendorf tubes containing the cells were centrifuged for 5 minutes at 13,500 rpm. Cells were washed twice with 3% bovine serum albumin in PBS after each fixation, permeabilization, and staining steps of the protocol. Cells were permeabilized with 1% saponin in PBS for 15 minutes at room temperature. Cells were stained for 30 minutes at room temperature with the EdU reaction mixture defined by the manufacturer’s protocol. Cells were stained for 15 minutes at room temperature with 1 μg/mL Hoechst nuclear stain in PBS. Finally, cells were resuspended in 30 μL PBS and stored at 4°C in the dark until imaging.

Prior to imaging, glass-slides of the stained *N. fowleri* were prepared. Slides were imaged on the Zeiss Elyra S1 super resolution microscope at 100x.

### Development of *in vitro* EdU drug susceptibility assay

Upon validation of EdU incorporation in *N. fowleri* trophozoites, we next wanted to develop a drug susceptibility assay that uses EdU as a measure of cell viability. EdU will only incorporate in cells that have undergone active DNA replication. We were interested in developing this assay to evaluate the ability of drugs, both current treatments and promising leads identified from high-throughput screens, to inhibit *N. fowleri* replication.

The Millipore ImageStream X MK II flow cytometer was used for EdU detection in drug-treated *N. fowleri* trophozoites. The ImageStream offers the tremendous advantage of phenotypic assessment of drug treatment on individual *N. fowleri* cells, while simultaneously obtaining quantitative data on EdU and Hoechst staining. The ImageStream flow cytometer requires a minimum of 1 x 10^6^ cells in 50 μL for detection. Based on the ImageStream requirements, a starting concentration of 20,000 *N. fowleri* trophozoites was used given that after 72 hours, trophozoites replicate to approximately 1.2 x 10^6^ cells. To optimize the ImageStream analysis protocol, 20,000 *N. fowleri* trophozoites were seeded in 500 μL NCM. 10 μM EdU was added at 48 hours. At 72 hours, trophozoites were fixed, permeabilized, and stained with the EdU reaction mixture and 1 μg/mL Hoechst as described above. Eppendorf tubes containing stained trophozoites in 30 μL PBS were taken to the Cytometry Shared Resource Laboratory for flow cytometry analysis.

An acquisition template was created on the Millipore ImageStream. A diameter gate of 7.5 μm – 30 μm was set. Images were acquired using the 375 nm and 488 nm lasers, as well as the brightfield lens. 10,000 events per tube were collected for analysis by the ImageStream. Events were collected using the 40x objective. Analysis of ImageStream data was performed on the IDEAS software. A fluorescence compensation matrix was created using single-color control tubes. A data analysis template was generated to examine compensated data files. Cell populations were gated based upon presence or absence of EdU incorporation.

### *In vitro* EdU drug susceptibility assay to assess current PAM therapeutics

Upon validation of EdU incorporation in *N. fowleri* trophozoites and development of a detection method using the Millipore ImageStream, current PAM therapeutics were assessed for their ability to inhibit *N. fowleri* replication. Azithromycin, amphotericin B, and miltefosine, were selected for assessment. Posaconazole was also selected as it has demonstrated promise as a potential novel PAM treatment [9]. All compounds were prepared as 5 mg/mL solutions in dimethyl sulfoxide (DMSO). Compounds were serially diluted to yield a concentration range of either 100 nM – 0.1 pM (posaconazole), 1 μM – 1 pM (amphotericin B, azithromycin), or 100 μM – 100 nM (miltefosine). Experimental Eppendorf tubes contained a total volume of 500 μL, consisting of 20,000 *N. fowleri* trophozoites and the appropriate drug concentration in NCM. Tubes were incubated at 34°C. At 48 hours, EdU was added to yield a concentration of 10 μM in the tube. At 72 hours, tubes were fixed, permeabilized, and stained with the EdU reaction mixture and 1 μg/mL Hoechst as described above. Single-color control tubes (EdU only, Hoechst only, no stain) were included for each replicate.

Labelled amoebae in tubes were analyzed on the Millipore ImageStream as described above. We collected data until 10,000 events were acquired or up to 5 minutes acquisition time. A 5-minute cut-off time was implemented due to tubes with higher concentrations of drug not having enough cells fitting the diameter gate to be counted. An acquisition time of 5 minutes was more than enough for 10,000 events to be counted in both control and drug-treated tubes. ImageStream data was analyzed by using the IDEAS software. For each drug series replicate, EdU positive and negative gates were created using the single-color controls from that run. All data analysis files were compensated. Using the percent of positive EdU cells in each tube, IC_50_ curves could be generated.

### Drug Susceptibility with CellTiter-Glo

Drug susceptibility assays using CellTiter-Glo 2.0 (Promega) were performed as previously described [6, 7, 9]. Azithromycin, amphotericin B, miltefosine, and posaconazole were evaluated in the CellTiter-Glo assay for comparison with IC_50s_ generated by the EdU assay. Briefly, all compounds were prepared as 5 mg/mL solutions in dimethyl sulfoxide (DMSO). Compounds were serially diluted to yield a concentration range of either 100 nM – 0.1 pM (posaconazole), 1 μM – 1 pM (amphotericin B, azithromycin), or 100 μM – 100 nM (miltefosine). Compounds were plated in white 96-well microtiter plates with 4,000 N. fowleri trophozoites and a total well volume of 100 μL. Plates were incubated at 34°C for 72 hours. At 72 hours, CellTiter-Glo reagent was added and luminescence was measured on the SpectraMax i3x (Molecular Devices) plate reader at 490 nm. IC_50s_ were generated using CDD Vault.

### Evaluation of EdA incorporation

EdA (7-Deaza-2′-deoxy-7-ethynyladenosine) is an adenosine analog that functions in a similar manner as the EdU molecule. EdA was assessed for its ability to incorporate in to DNA of *N. fowleri* trophozoites during replication.

A similar protocol as was described above for investigation of EdU incorporation was used for EdA. In short, 20,000 trophozoites were seeded with 500 μL NCM and 10 μM EdA in Eppendorf 1.5 mL centrifuge tubes for 72 hours and incubated at 34°C. At 72 hours, the manufacturer’s protocol for EdU detection was followed with minor modifications. Cells were stained with the EdU reaction mixture and 1 μg/mL Hoechst nuclear stain. Cells were assessed for EdA incorporation using the previously described Millipore ImageStream analysis protocol.

## Acknowledgements

We thank Julie Nelson and the University of Georgia Cytometry Shared Resource Laboratory for excellent assistance and support with the Millipore ImageStream. We also thank Dr. Muthugapatti Kandasamy and the University of Georgia Biomedical Microscopy Core for training and use of the Zeiss ELYRA S1 microscope. Funding for this study was provided in part by NIH grants R03AI149097 and S10OD021719.

